# Over 1000 tools reveal trends in the single-cell RNA-seq analysis landscape

**DOI:** 10.1101/2021.08.13.456196

**Authors:** Luke Zappia, Fabian J. Theis

## Abstract

Recent years have seen a revolution in single-cell technologies, particularly single-cell RNA-sequencing (scRNA-seq). As the number, size and complexity of scRNA-seq datasets continue to increase, so does the number of computational methods and software tools for extracting meaning from them. Since 2016 the scRNA-tools database has catalogued software tools for analysing scRNA-seq data. With the number of tools in the database passing 1000, we take this opportunity to provide an update on the state of the project and the field. Analysis of five years of analysis tool tracking data clearly shows the evolution of the field, and that the focus of developers has moved from ordering cells on continuous trajectories to integrating multiple samples and making use of reference datasets. We also find evidence that open science practices reward developers with increased recognition and help accelerate the field.

## Introduction

Developments in single-cell technologies over the last decade have drastically changed the way we study biology. From measuring genome-wide gene expression in a few cells in 2009 [1], researchers are now able to investigate multiple modalities in thousands to millions of cells across tissues, individuals, species, time and conditions [2,3]. Commercialisation of these techniques has improved their robustness and made them available to a greater number of biological researchers. Although single-cell technologies have now extended to other modalities including chromatin accessibility [4,5], DNA methylation [6,7], protein abundance [8] and spatial location [9,10], much of the focus of the single-cell revolution has been on single-cell RNA sequencing (scRNA-seq). Single-cell gene expression measurements are cell type-specific (unlike DNA), more easily interpretable (compared to epigenetic modalities) and scalable to thousands of features (unlike antibody-based protein measurements) and thousands of cells. These features mean that scRNA-seq can be used as an anchor, often measured in parallel and used to link other modalities.

While single-cell assays of all kinds are now more readily available, the ability to extract meaning from them ultimately depends on the quality of computational and statistical analysis. With the rise of new technologies we have seen a corresponding boom in the development of analytic methods. After years of rapid growth the sheer number of possible analysis options now available can be bewildering to researchers faced with an scRNA-seq dataset for the first time. Efforts have also been made to benchmark common tasks such as clustering of similar cells [11,12], differential expression between cells [13,14] or integration of multiple samples [15,16] in an attempt to establish which approaches are consistently good performers and in which situations they fail. Building on these benchmarks the community has now produced tutorials [17], workshops and best practices recommendations for approaching a standard analysis [18,19].

Several projects exist which attempt to chart the progression of scRNA-seq technologies, datasets and analysis tools. For example, the Single Cell Studies database tracks the availability and size of scRNA-seq datasets and has been used to show trends in technologies and analysis [20]. The Awesome Single Cell repository is a community-curated list of software packages, resources, researchers and publications for various single-cell technologies [21] and Albert Villela’s SingleCell Omics spreadsheet [22] tracks a range of information including technologies, companies and software tools. While these are all very useful resources they either have different focuses or are less structured and detailed than the scRNA-tools database.

The scRNA-tools database focuses specifically on the cataloguing and manual curation of software tools for analysing scRNA-seq data [23]. When tools become available (usually through a bioRxiv preprint) we classify them according to the analysis tasks they can be used for and record information such as associated preprints and publications, software licenses, code location, software repositories and a short description. All this information is publicly available in an interactive format at https://www.scrna-tools.org/. As the number of tools in the database has moved past 1000 we have taken this opportunity to provide an update on the current state of the database, explore trends in scRNA-seq analysis across the past five years. We find that the focus of tool developers has moved on from continuous ordering of cells to methods for integrating samples and classifying cells. The database also shows us that more new tools are built using Python while the relative usage of R is declining. We also examine the role of open science in the development of the field and find that open source practices lead to increased citations. While the scRNA-tools database does not record every scRNA-seq analysis tool the large proportion it does include over the history of what is still a young field make these analyses possible and a reasonable estimate of trends across all tools.

## Results

### Current state of scRNA-seq analysis tools

We first started cataloging software tools for analysing scRNA-seq data in 2016 and the scRNA-tools database currently contains 1027 tools as of 12 August 2021 (Figure 1A). This represents a more than tripling of the number of available tools since the database was first published in June 2018. The continued growth of the number of available tools reflects the growth in the availability of and interest in single-cell technologies. It also demonstrates the continued need for new methods to extract meaning from them. This trend has continued for more than five years and if it continues at the current rate we can expect to see around 1500 tools by the end of 2022 and more than 3000 by the end of 2025.

**Figure 1:**
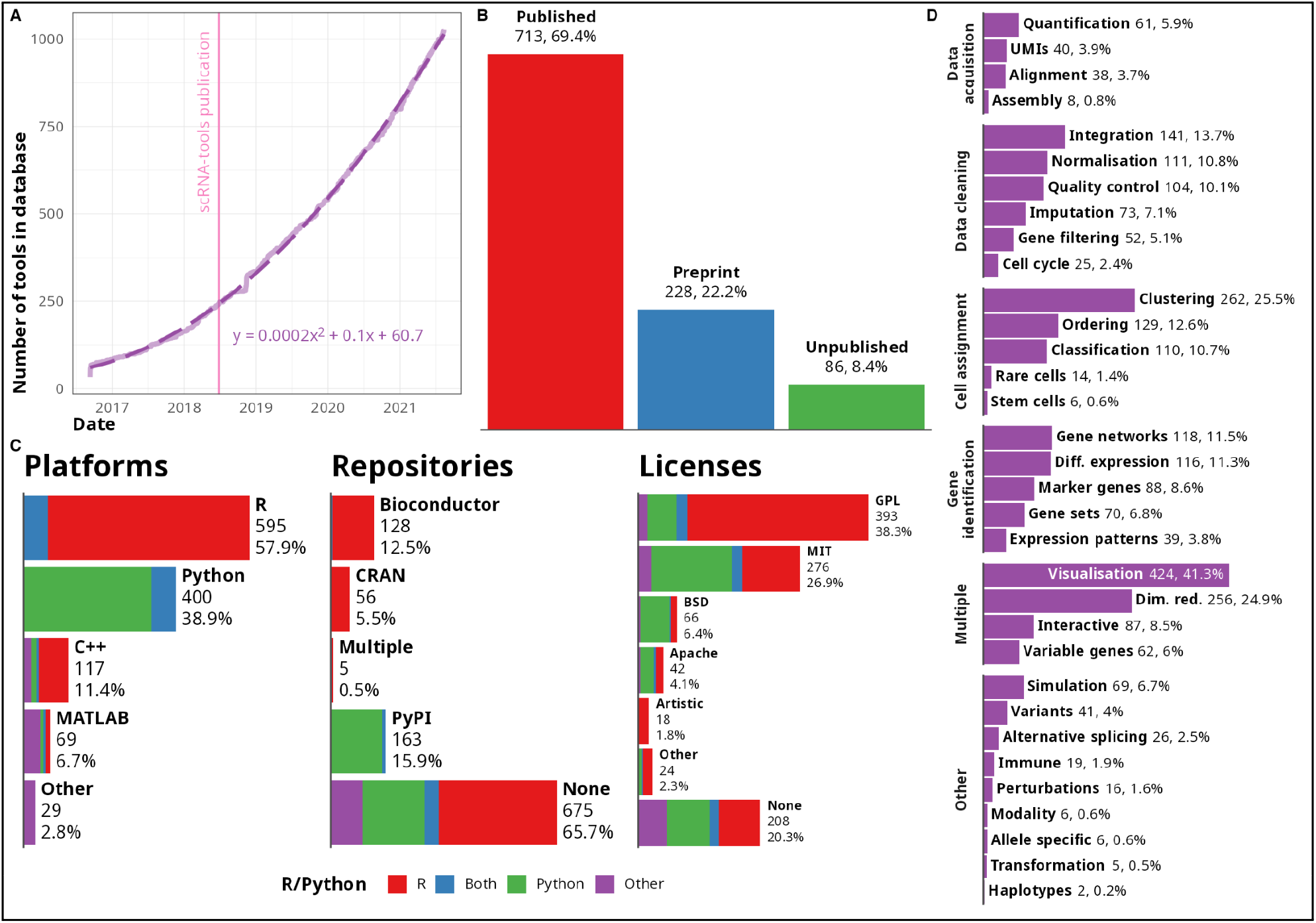
Overview of the scRNA-tools database. A) Line plot of the number of tools in the scRNA-tools database over time. The development of tools for analysing scRNA-seq data has continued to accelerate with more than 1000 tools currently recorded. Dotted line shows a quadratic fit (*y* = 0.0002*x*^2^ +0.1*x* + 60. 7). B) Publication status of tools in the scRNA-tools database. Around 70 percent of tools have at least one peer-reviewed publication while more than 20 percent have an associated preprint. C) Bar charts showing the distribution of platforms, software licenses and software repositories for tools in the scRNA-tools database. Colours indicate proportions of tools using R or Python. D) Bar chart showing the proportion of tools in the database assigned to each analysis category. Categories are grouped by broad phases of a standard scRNA-seq analysis workflow.

#### Publication status

The scRNA-tools database records preprints and publications associated with each tool, currently including more than 1400 references. Around two thirds of tools have at least one peer-reviewed publication while another quarter have been described in a preprint but are not yet peer-reviewed (Figure 1B). The remaining 10 percent currently have no associated references and have come to our attention through other means such as Twitter, software and code repositories or submissions via the scRNA-tools website. The overall number of tools with peer-reviewed publications has increased over time, which is to be expected as tools make their way through the publication process. For more discussion of the delay in publication and the effect of preprints see the open science section.

#### Software platforms, licenses and repositories

Tool developers must make a choice of which platform to use and this is also recorded in the scRNA-tools database. In most cases the platform is a programming language but a minority of tools are built around an existing framework including workflow managers such as Snakemake [24,25] and Nextflow [26]. R [27] and Python [28] continue to be the dominant platforms for scRNA-seq analysis, just as they are for more general data science applications (Figure 1C). C++ continues to play a role, particularly for including compiled code in R packages to improve computational efficiency. Although it is a minority (around seven percent) there is a consistent community of developers focused on MATLAB. The popularity of interpreted languages (R, Python, MATLAB) with commonly used interactive interfaces rather than compiled languages (C++) is consistent with relatively few tools being developed for low-level, computationally-intensive tasks such as alignment and quantification where compiled languages are more commonly seen (Supplementary Figure 1).

Around two thirds of tools are not available from a standard centralised software repository (CRAN, Bioconductor or PyPI) and are mostly available only from GitHub. While this makes it harder for other members of the community to install and use these tools it is perhaps unsurprising given the large amount of time and effort required to maintain a software package. Many of these tools may also be primarily intended as example implementations of a method rather than a tool designed for reuse by the community. A higher proportion of Python-based tools are available from PyPI (the primary Python package repository) when compared to those built using R. This may reflect the lower submission requirements of PyPI which does not enforce checks for documentation or testing unlike R repositories. Of R packages available from central repositories the majority of developers have chosen to submit their tools to the biology-focused Bioconductor [29] repository rather than the more general CRAN. The Bioconductor community is well-established and provides centralised infrastructure (such as the commonly used SingleCellExperiment class) designed to allow small, specialised packages to work together.

Most tools are covered by a standard open-source software license, although there remains a consistent minority of tools (around 20 percent) for which no clear software license is available. The lack of a license can severely restrict how tools can be used and the ability of the community to learn from and extend existing code. We strongly encourage authors to clearly license their code and for reviewers and journal editors to include checks for a software license into the peer-review process to avoid this problem. Among those tools that do have a license, variants of the copy-left GNU Public License (GPL) are most common (particularly amongst R-based tools). The MIT license is also used by many R tools but is more common for Python tools, as are BSD-like and Apache licenses. These licenses all allow reuse of code but may impose some conditions such as retaining the original copyright notice [30]. GPL licenses also require that any derivatives of the original work are also covered by a GPL license.

#### Analysis categories

A unique feature of the scRNA-tools database is the classification of tools according to which analysis tasks they can be used for. These categories have been designed to capture steps in a standard scRNA-seq workflow and as new tasks emerge additional categories can be created. While they have limitations, these categories should provide some guidance to analysts looking to complete a particular task and can be used to filter tools on the scRNA-tools website. Categories that are applicable to many stages of analysis (visualisation, dimensionality reduction) are among the most common, as is clustering which has been the focus of much tool development but is also required as an input to many other tasks (Figure 1D). Other stages of a standard analysis form the next most common categories including integration of multiple samples, batches or modalities, ordering of cells into a lineage or pseudotime trajectory, quality control of cells, normalisation, classification of cell types or states and differential expression testing. Tools that either construct or make use of gene networks are also common. Some tools such as Seurat [31] and Scanpy [32] are general analysis tool boxes which can complete many tasks, while others are more specialised and focus on one problem. While the number of categories per tool is variable there is no clear trend in tools becoming more general or specialised over time (Supplementary Figure 2).

Other categories in the scRNA-tools database capture some of the long tail of possible scRNA-seq analyses. For example, analysis of alternative splicing or allele-specific expression, stem cells, rare cell types and immune receptors may not be relevant for all experiments but when they are having methods for those specific tasks can be invaluable. As biologists use scRNA-seq to investigate more phenomena, developers will create methods and tools for more specific tasks. We plan to update the categories in the scRNA-tools database to reflect this and have recently added a category for tools designed to work with perturbed data such as drug screens or gene editing experiments including MELD [33], scTenifoldKnK [34] and scGen [35].

### Trends in scRNA-seq analysis tools

Over five years of data in the scRNA-tools database on new tools and their associated publications allows us to track some of the trends in scRNA-seq analysis over that time. Here we focus on trends in analysis tasks as well as choice of development platform.

#### Increasing proportion of tools use Python

Figure 2A shows how the proportion of tools using the most common programming languages has changed over time as more tools are added to the database. The clear trend here is the increasing popularity of Python and the corresponding decrease in the proportion of tools built using R. There are several possible explanations for this trend.

**Figure 2:**
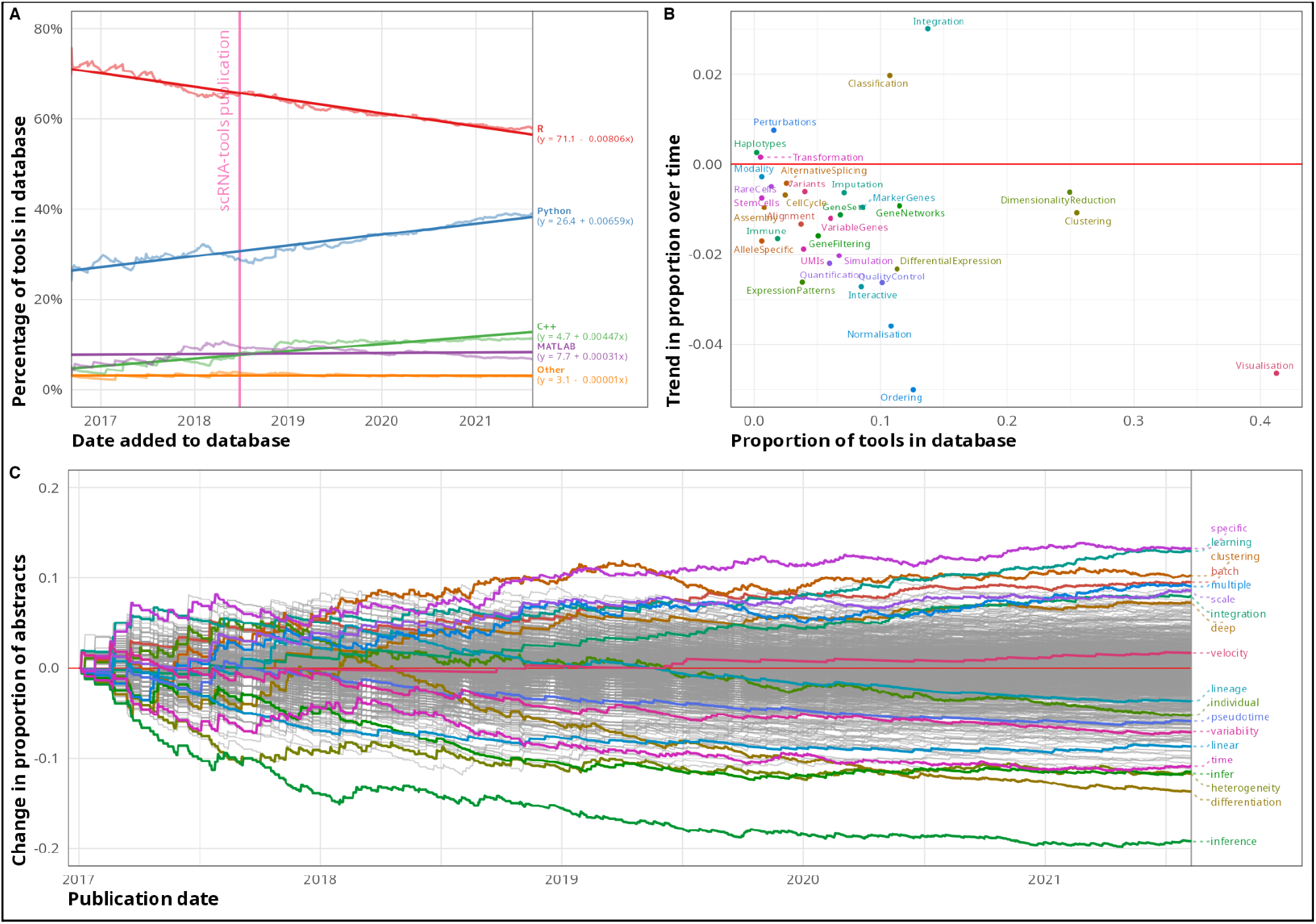
Trends in scRNA-seq analysis tools. A) Line plot of platform usage of tools in the scRNA-tools database over time. Python usage has increased over time while R usage has decreased. Darker lines show linear fits with coefficients given in the legend. Vertical pink line shows the date of the original scRNA-tools publication. B) Scatter plot of trends in scRNA-tools analysis categories over time. Current proportion of tools in the database shown on the x-axis and trend in proportion change shown on the y-axis. C) Line plot of trend in word use in scRNA-seq analysis tools publication abstracts over time. Publication date is shown on the x-axis and change in proportion of abstracts containing a word on the y-axis. Some important and highly variable terms are highlighted.

As the size and complexity of scRNA-seq datasets has increased, the potential memory and computational efficiency of Python has become more relevant. Another possible catalyst is the development of Python-based infrastructure for the community to build around such as the AnnData and Scanpy packages [32] which play a similar central role in the Python environment as the SingleCellExperiment and Seurat packages do in the R environment (Supplementary Figure 3). These standard representations improve interoperability between packages and allow developers to focus on analysis methods rather than how to store their data which may have previously been a barrier for Python developers.

While bulk transcriptomics typically focused on statistical analysis of a designed experiment, scRNA-seq analysis is often more exploratory and employs more machine learning techniques such as unsupervised clustering and more recently various neural-network architectures. This shift in analysis focus may have triggered a corresponding shift in demographics with more researchers from a computer science background turning their attention to developing scRNA-seq analysis methods and bringing with them a preference for Python over R. If this trend continues at the current rate we can expect Python to overtake R as the most common platform for scRNA-seq analysis tools by mid 2025, however R will continue to be an important platform for the community. It is also important to note that this trend represents the preferences of developers and may not reflect how commonly these platforms are used by analysts.

#### Greater focus on integration and classification

Trends also exist in the tasks that new tools perform. In Figure 2B we can see the overall proportion of tools in the database assigned to each category against the trend in proportion over time. Two categories stand out as increasing in focus over time: integration and classification. Both of these trends reflect the growing scale, complexity and availability of scRNA-seq datasets. While early scRNA-seq experiments usually consisted of a single sample or a few samples from a single lab it is now common to see experiments with multiple replicates, conditions and sources. For example studies have benchmarked single-cell protocol across multiple centres [36], measured hundreds of cell lines at multiple time points [37] and compared immune cells between cancer types [38]. Handling batch effects between samples is vital to producing meaningful results but can be extremely challenging due to the balance of removing technical effects while preserving biological variation. The importance of this task is demonstrated by the more than 130 tools that perform some kind of integration and is particularly relevant for global atlas building projects like the Human Cell Atlas [39] which attempt to bring together researchers and samples from around the world to map cell types in whole tissues or organisms. Recent technological advances have made it more feasible to measure biological signals other than gene expression in individual cells. Some tools tackle this more challenging task, either using one modality to inform analysis of another or by bringing modalities together for a fully integrated analysis. Combining multiple data types can provide additional insight by confirming a signal that is unclear in one modality (for example protein expression confirming gene expression) or revealing another aspect of a biological process (chromatin accessibility used to show how genes are regulated).

The increased interest in classification can also be seen as an attempt to tackle the increasing scale of scRNA-seq data. Rather than performing the computationally and labour intensive task of merging datasets and jointly analysing them to get consistent labels, classifier tools make use of public references to directly label cells with cell types or states. This approach is a shortcut for analysts, allowing them to skip many early analysis steps but is limited by the completeness and reliability of the reference. For this reason integration and classification are intimately linked, with effective integration required to produce quality references for classification. Some tools address both sides of this problem, integrating datasets to produce a reference and providing methods to classify new query datasets while considering batch effects in the query. Once more comprehensive references atlases are available this reference-based workflow will likely replace the current unsupervised clustering approach for many analyses.

#### Decrease in ordering and common tasks

The category with the biggest decreasing trend over time is the ordering category, which refers to tools which determine a continuous order for cells, usually related to a developmental process or another perturbation such as onset of disease or effect of a drug treatment. This category represents perhaps the biggest promise of the single-cell revolution, the ability to interrogate continuous biological processes at the level of individual cells. Some of the earliest scRNA-seq analysis tools addressed this task but the proportion of new tools containing ordering methods has significantly decreased since. It is unclear why this is the case. It may simply be that high-performing methods have been established [40] and adopted by the community, reducing the need for further development. Alternatively it could be that the initial excitement was overcome by limitations revealed when these techniques were applied to real datasets [41]. A subset of the ordering category are RNA velocity methods [42,43] which offer another approach to analysing continuous processes but has resulted in the development of relatively few new tools.

Many of the other categories show some trend towards decreasing proportions with normalisation and visualisation having the biggest reductions. A plausible explanation for these changes is the consolidation of common tasks into analysis tool boxes with each of the major software repositories having standard workflows based around a few core tools. With these available and accessible through the use of standard data structures developers no longer need to implement each stage of an analysis and can focus on particular tasks.

#### Trends in publication abstracts

Similar trends can be seen in the text of publications associated with analysis tools. Figure 2C shows how the proportion of abstracts associated with scRNA-seq tools containing keywords has changed since 2017. Highly variable words are highlighted as well as some related to trends discussed above. Both “batch” and “integration” have become more common, mirroring the increase in tools performing integration, as have related terms like “scale” and “multiple”. Machine learning terms “deep” and “learning” have also become more common, consistent with the increased use of Python which is the primary language for deep-learning. In contrast “lineage”, “pseudotime” and “differentiation” have all decreased consistent with the reduction of tools in the ordering category. The “velocity” term has seen a small increase in use over the last two years but as these abstracts only come from publications associated with tools (and not publications that focus on analysis of scRNA-seq data) it is difficult to say anything about the take-up of these methods in the community and whether they have replace the earlier generation of ordering techniques.

### Open science accelerates scRNA-seq tool development

Researchers must make a conscious choice about when and how to share their work and for developers of software tools there are several options. A tool could be made available during development, when there is a stable version ready for users, accompanying a preprint or only after a peer-reviewed publication. Here we touch on the decision around open science practices and the effect they have on scRNA-seq analysis tools.

#### GitHub is the primary home for scRNA-seq analysis tools

The vast majority of tools in the scRNA-tools database (over 90 percent) have a presence on the social coding website GitHub. Like GitLab, BitBucket and other similar services GitHub provides an all-in-one service for open-source software development which has been embraced by the scRNA-seq community. Being available on GitHub allows the community to ask questions, raise issues, suggest enhancements and contribute features. Across the scRNA-tools database there are 960 associated GitHub repositories from 709 owners (Figure 3A). To these repositories over 1700 contributors have made more than 150000 commits and opened almost 28000 thousand issues. If each of these commits and issues represents just 10 minutes of work on average that corresponds to more than 31000 person hours of work or three and half person years. This is a tremendous amount of effort from the community but is likely still a large underestimate as this doesn’t capture many of the tasks involved in software development and maintenance.

**Figure 3:**
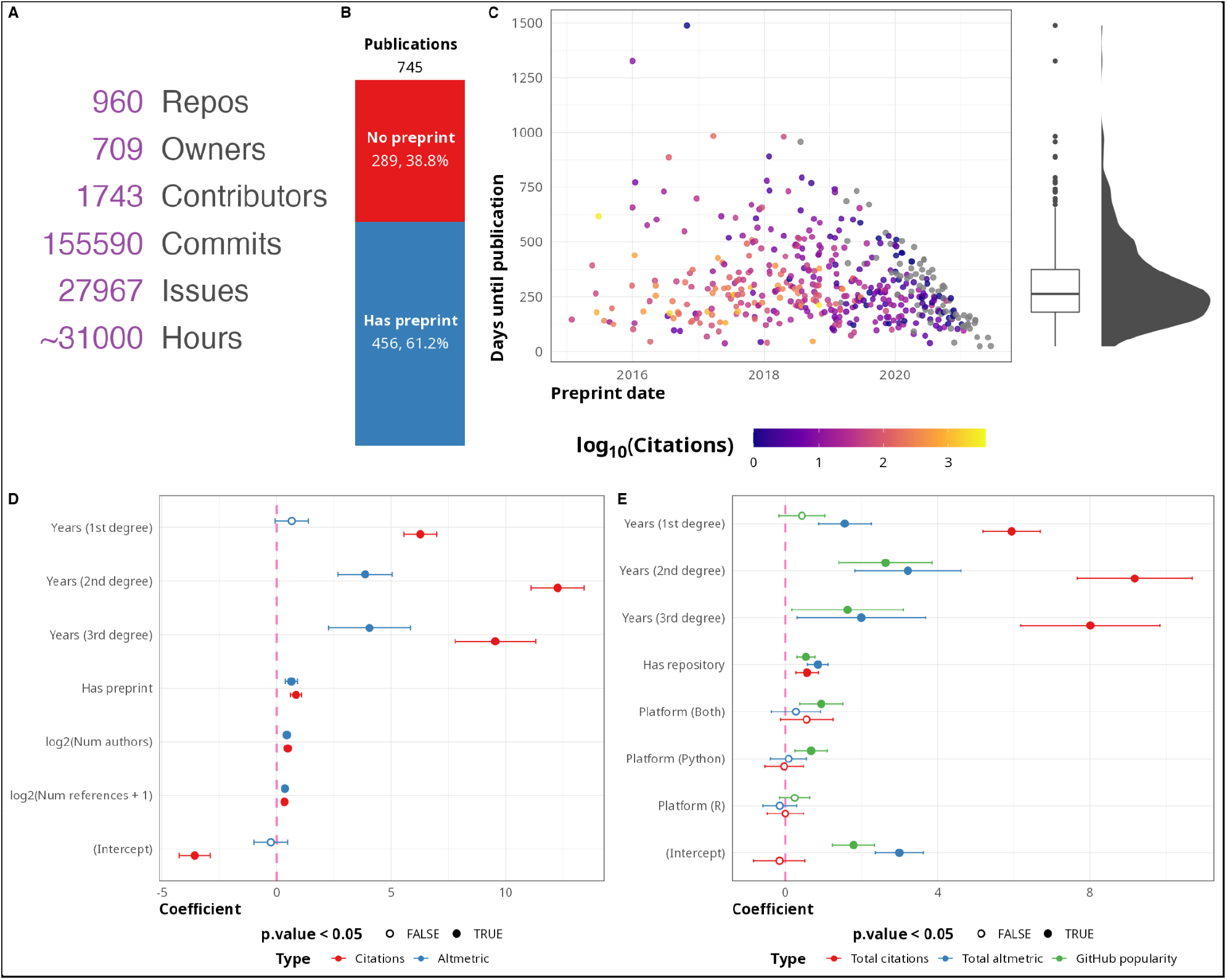
Open science in scRNA-seq tools development. A) GitHub summary statistics for scRNA-seq tool repositories. B) Stacked bar plot showing the proportion of publications with and without an associated publication. C) Scatter plot showing preprint date against number of days until publication, colours indicate number of citations (log scale). Box plot and density on the right show the distribution of time delay in publication. D) Coefficients for linear models predicting citations and Altmetric score for publications. Years since publication is modelled as a cubic spline with three degrees of freedom. Error bars show a 95 percent confidence interval. E) Coefficients for linear models predicting total citations, total Altmetric score and GitHub popularity score for tools. Error bars show a 95 percent confidence interval.

#### Preprints allow the rapid development of scRNA-seq methods

Around 60 percent of the 745 publications associated with tools in the scRNA-tools database were preceded by a preprint (Figure 3B). Figure 3C shows the number of days between preprints and publication of the final peer-reviewed article. The average delay in publication is around 250 days but the biggest gap is around six times that long at almost 1500 days. The willingness of the scRNA-seq community to share their work in preprints and make code implementing it on GitHub is a big contributor to the rapid development of the field. Without early sharing of ideas we would still be waiting for tools that were released a year ago and that delay would likely be much longer if we consider the compounding effect of early access over time.

#### Open science practices lead to increased citations

To quantify the effect of open science practices such as posting preprints and sharing code we modelled citations using a method based on that suggested by Fu and Hughey [44]. We acknowledge that citations are a flawed and limited metric for assessing software tools, for example they only indirectly measure the effectiveness and usability of a tool. Despite these flaws, citations are still the metric most commonly used to judge researchers and their work so it is important to see how they are affected by open science practices. The same model is also used to predict Altmetric score and GitHub popularity (for tools).

Figure 3D shows coefficients for a log-linear model predicting metrics for publications. Unsurprisingly the biggest predictor of number of citations is years since publication (included in the model as a splice with three degrees of freedom) (Supplementary Table 1). Whether or not a publication has an associated preprint was also a significant positive predictor (coefficient = 0.85, 95% CI = (0.60, 1.1), p-value = 3.1e-11). As we modelled metrics on a log_2_ scale a coefficient close to one indicates a two-fold increase in citations for publications with a preprint. The number of authors and number of references were also significant in this model but with smaller effect sizes. Results for Altmetric are broadly similar to those seen for citations. This same effect was observed by Fu and Hughey more generally across fields and as well as other studies [45]. That preprints both help the community and result in more citations should encourage more researchers to share their work in this way and outweigh the fear of being “scooped”.

Coefficients for a similar model for predicting metrics at the tool level are shown in Figure 3E. In this model we replaced having a preprint with whether or not the tool is available from one of the major software repositories (CRAN, Bioconductor, PyPI). We also included programming language as a possible predictor. As well as citations and Altmetric score for tools we also measured “GitHub popularity”, a score combining number of GitHub stars and forks. Although the effect is smaller, publishing tools in a software repository also had a significant effect on all three metrics (Supplementary Table 2). This makes sense as tools available from repositories are complete packages which should be easier for users to install and therefore lead to more citations. Programming language was not significant for predicting citations or Altmetric but did have a positive effect for Python tools in predicting GitHub popularity.

## Conclusion

Interest in single-cell RNA-sequencing and other technologies continues to increase with more datasets being produced with more complex designs, greater numbers of cells and multiple modalities. To keep pace with the increase in datasets there has been a corresponding increase in the development of computational methods and software tools to help make sense of them. We have catalogued this development in the scRNA-tools database which now records more than 1000 tools. While we try to record every new scRNA-seq analysis tool the database is likely incomplete. We warmly welcome contributions to the database and encourage the community to submit new tools or updates via the scRNA-tools website.

Despite the incompleteness of the database it is a large sample of the scRNA-seq analysis landscape and using it we can observe trends in the field. We see that there has been a decrease in development of methods for ordering cells into continuous trajectories and an increased focus on methods for integrating multiple datasets and using public reference datasets to directly classify cells. The classification of tools by analysis task is an important feature of the scRNA-tools database and we plan to expand these categories to cover more aspects of scRNA-seq analysis. We also examined trends in development platforms and found that more new tools are being built using Python than R. While it is exciting to see the field develop, the continued increase in the number of tools presents some concerns. There is still a need for new tools but if growth continues at this rate it is a risk that the community will start repeating work and approaches that are already available. By providing a catalogue of tools in a publicly available website the scRNA-tools database makes it easier to find current tools and we encourage developers to contribute to existing projects where that is a good fit. Equally important are continued, high-quality benchmarking studies to rigorously evaluate the performance of methods and we hope to include this information in future versions of the database. This would include published benchmarks but also results from continuous benchmarking efforts such as those proposed by the Open Problems in Single-Cell Analysis project (https://openproblems.bio/).

The scRNA-seq community has largely embraced open science practices and we sought to quantify their effect on the field. We found an average delay of 250 days between preprints and peer-reviewed publications with some examples being much longer. The willingness of researchers to share early versions of their work has likely been a major contributor to the rapid development of the field. We also found that open science did not conflict with recognition of work, with both preprints and availability of tools in software package repositories being a positive predictor of citations and Almetric scores.

We hope that the scRNA-tools database is a valuable resource for the community, both for helping analysts find tools for a particular task and tracking the development of the field over time.

## Methods

### Curation of the scRNA-tools database

The main sources of tools and updates for the scRNA-tools database are Google scholar and bioRxiv alerts for scRNA-seq specific terms (Table 1). Other sources include social media, additions to similar projects like the Awesome Single Cell page [21] and submissions via the scRNA-tools website. Once a potential new tool is found it is checked if it fulfills the criteria for inclusion in the database, namely that it can be used to analyse scRNA-seq data and that the tool is available to users to install and run locally (this excludes tools and resources that are only available online). Most of the information for new tools (description, license, categories) comes from code repositories (GitHub) and package documentation.

**Table 1:** Alert terms for Google Scholar and bioRxiv

| Google Scholar | bioRxiv |
| --- | --- |
| "scRNAseq" OR "scRNA-seq" OR "sc-RNA-seq" OR "(sc)RNA-seq" | scRNA-seq scRNAseq |
| "single-cell gene expression" | single-cell RNA-sequencing |
| "single-cell transcriptomics" OR "single-cell transcriptome" |  |
| "single-cell RNA sequencing" |  |
| "single-cell rna-seq" |  |

A command line application written in R (v4.0.5) [27] and included in the main scRNA-tools repository is used to make changes to the database. Using this interface rather than editing files allows input to be checked and some information to be automatically retrieved from Crossref (using the *rcrossref* package (v1.1.0) [46]) and GitHub (using the *gh* package (v1.2.0) [47]). This application also contains functionality for performing various consistency checks including identifying new or deprecated software package and code repositories, updated licenses and new publications linked to preprints. Identification of new software package repositories is done by fuzzy matching of the tool name using the *stringdist* package (v0.9.6.3) [48]. Linking of preprints uses an R implementation of the algorithm by Cabanac, Oikonomidi and Boutron [49] available in the *doilinker* package (v0.1.0) [50]. Software licenses are standardised using the SPDX License List [51]. Data files and visualisations shown on the scRNA-tools website are also produced by the command line application using *ggplot2* (v3.3.3) [52] and *plotly* (v4.9.2.2) [53]. Dependencies for the application and managed using the *renv* package (v0.13.0) [54] and more details on its use and functionality can be found at https://github.com/scRNA-tools/scRNA-tools/wiki.

### Contributing to scRNA-tools

Contributions to the scRNA-tools database from the community are welcomed and encouraged. The easiest way to contribute is by submitting a new tool or update using the form on the scRNA-tools website (https://www.scrna-tools.org/submit). For those comfortable using Git and GitHub changes can be made directly to the database and submitted as a pull request. Suggestions for changes or enhancements to the website can be made by opening issues on the scRNA-tools GitHub repository (https://github.com/scRNA-tools/scRNA-tools) or using the contact form on the website (https://www.scrna-tools.org/contact).

### Data acquisition and analysis

The main source of data for the analysis presented here is the scRNA-tools database as of 12 August 2021. This was read into R directly from the GitHub repository using the *readr* package (v1.4.0) [55] and manipulated using other *tidyverse* (v1.3.1) packages [56], particularly *dplyr* (v1.0.6) [57], *tidyr* (v1.1.3) [58], *forcats* (v0.5.1) [59] and *purrr* (v0.3.4) [60]. Additional information about references was obtained from the Crossref and Altmetric APIs using the *rcrossref* (v1.1.0) [46] and *rAltmetric* (v0.7.0) [61] packages. Dates for publications can vary depending on what information journals have submitted to Crossref. We used the online publication date where available, followed by the print publication date and the issued date. When a date was incomplete it was expanded to an exact date by setting missing days to first of the month and missing months to January, so that an incomplete date of 2021-08 would become 2021-08-01 and 2021 would become 2021-01-01. Dependencies for CRAN packages were obtained using the base R *package_dependencies* function and for Bioconductor packages using the *BiocPkgTools* package (v1.10.1) [62]. Python package dependencies were found using the Wheelodex API [63] and the *johnnydep* package (v1.8) [64]. All of the data acquisition and analysis was organised using a *targets* (v0.4.2) [65] pipeline with dependencies managed using *renv* (v0.13.2) [54].

#### Modelling trends

The trend in platform usage over time was simply modelled by calculating the proportion of tools using each of the main platforms on each day since the creation of the database. This proportion was then plotted over time to display the trend.

A more complex approach was taken to modelling the trend in categories. Time since the start of the database was divided into calendar quarters (three month periods) and for each category the proportion of tools added during each quarter was calculated. These quarter proportions were used as the input to a linear model (base R lm function) and the calculated slope taken as the trend for each category. These trends were then plotted against the overall proportion in the database to show the relationship between this and the trend over time.

Modelling of publication term usage over time started with abstracts obtained from Crossref. Abstracts were available for 1004 references. Each abstract was converted to a bag of words using the *tidytext* package (v0.3.1) [66] and URLs, numbers and common stop words were removed. We also excluded a short list of uninformative common scRNA-seq terms and those that appeared in less than 10 abstracts. For each word we calculated the cumulative proportion of abstracts that continued that word for each day since the first publication. The proportion as of the start of 2017 was taken as a baseline and the change since then plotted. A set of top words to display was chosen based on their variability over time, high absolute change in proportion and a few relevant to other parts of the analysis.

#### Modelling the effect of open science

To model the effect of a previous preprint on citations and Altmetric score for a publication we used a simplified version of the log-linear model proposed by Fu and Hughey [44]:

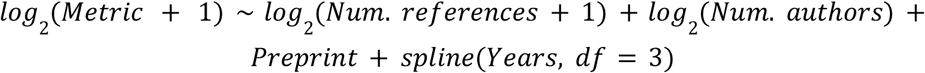

Here *Metric* is either citations or Altmetric score, *Preprint* is a boolean indicator variable showing whether or not the publication has an associated preprint and *spline*(*Years, df* = 3) is a natural cubic spline fit to years since publication with three degrees of freedom. We excluded additional author terms from the original model including whether an author had a US affiliation, whether an author had a Nature Index affiliation and the publication age of the last author. Information for these terms is difficult to collect and in the original publication they were shown to have a small effect compared to presence of a preprint. We also fit all publications together rather than for each journal individually.

We then adapted this model to consider tools rather than individual publications:

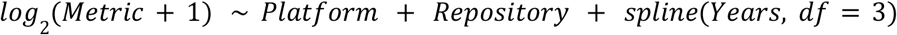

Here metric is either total citations for all publications and preprints associated with a tool, total Altmetric score for all publications and preprints or GitHub popularity score. GitHub popularity is calculated using number of forks of the repository and number of stars normalised by the age of the repository in years:

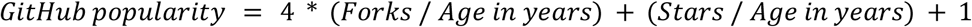

The *Platform* variable is an indicator showing whether the tool uses R, Python, both or some other platform (baseline). *Repository* is a boolean indicator showing whether the tool is available from any of the major software package repositories (CRAN, Bioconductor or PyPI).

Models were fit using the base R *lm* function and summary statistics including confidence intervals and p-values were extracted using the *ggstatsplot* package (v0.8.0) [67].

#### Visualisation

Plots and other figures were produced in R using the *ggplot2* package (v3.3.5) [52]. Various extension packages were also used including *ggtext* (v0.1.1) [68] for complex formatting of text and *ggrepel* (v0.9.1) [69] for labelling points and lines. Labels for bar charts and other plots were constructed using the *glue* package (v1.4.2) [70]. The final figures were assembled using the *cowplot* (v1.1.1) [71] and *patchwork* (v1.1.1) [72] packages.

## Availability

The scRNA-tools website and database is publicly available at https://www.scrna-tools.org/. The raw database files as well as code for managing the database and website is available on GitHub at https://github.com/scRNA-tools/scRNA-tools. Code and data files for the analysis presented here can be found on GitHub at https://github.com/scRNA-tools/1000-tools-paper.

## Acknowledgements

We would like to thank everyone in the community who has contributed to scRNA-tools since it began, including those who submitted new tools or updates, contributed code or provided advice and discussion. We also want to acknowledge the work of the developer community who continue to produce new tools and methods to drive the field forward. Lastly thank you to Fabiola Curion, Carlos Talavera-Lopez, Malte Lücken, Lukas Huemos and Karin Hrovatin for their commits on drafts of the manuscript.

## Conflict of interest

F.J.T. reports receiving consulting fees from Cellarity Inc., and ownership interest in Cellarity, Inc. and Dermagnostix.

## Supplementary information

**Supplementary Figure 1:**
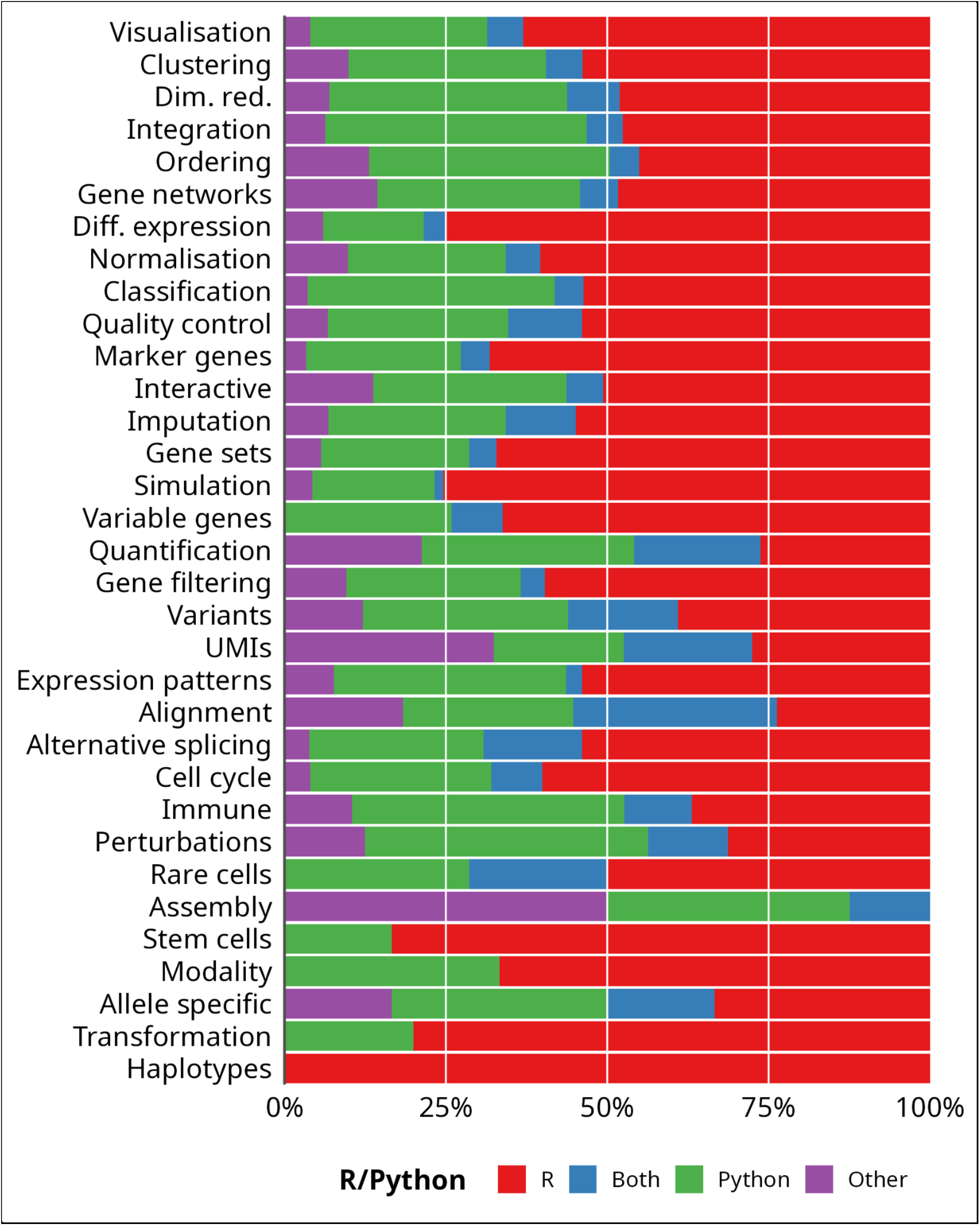
Stacked bar chart of proportions of tools in each analysis category built using R, Python or other platforms.

**Supplementary Figure 2:**
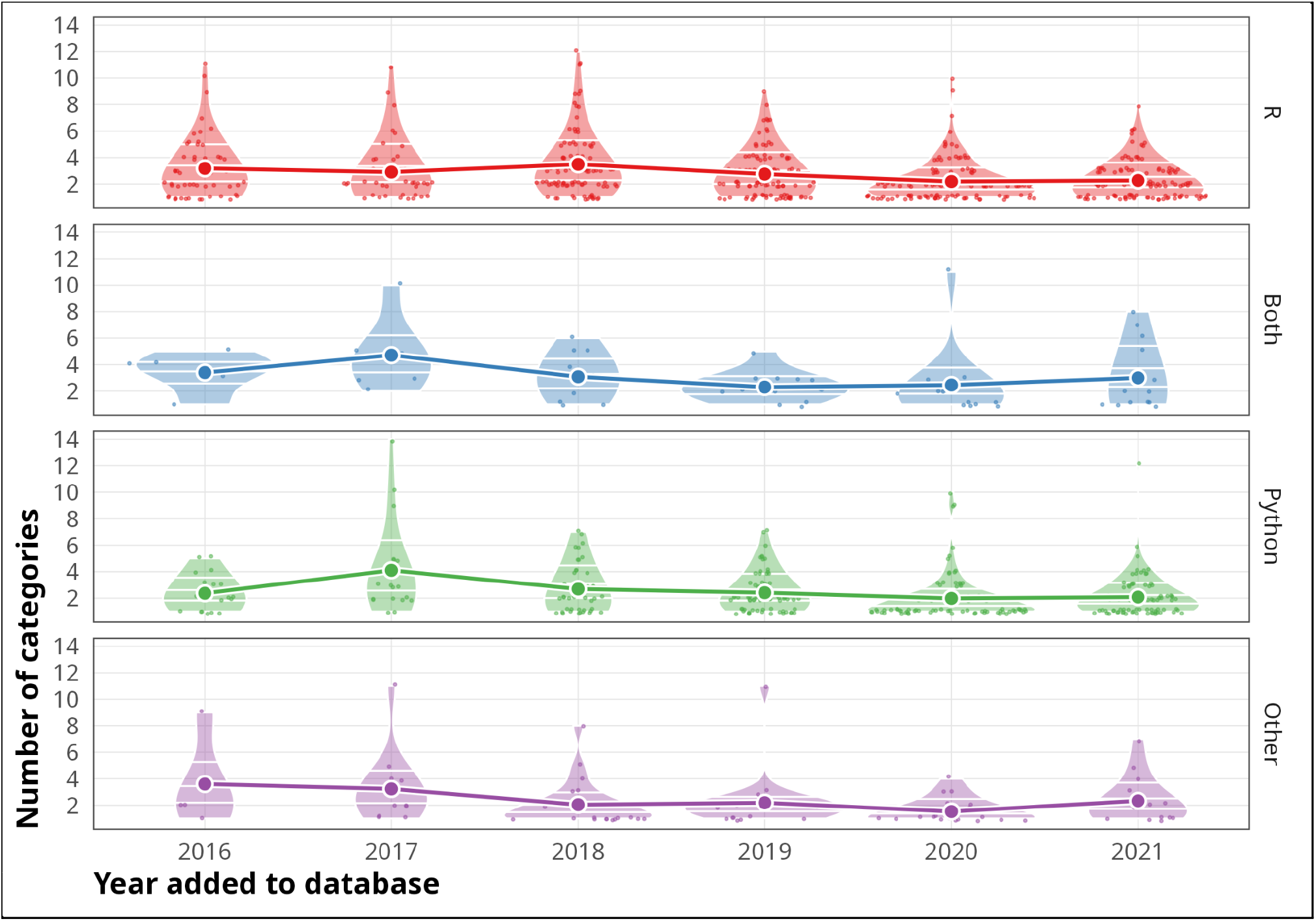
Number of categories per tool. Violin plots are shown divided by year and major platforms with points representing individual tools. Large points connected by lines indicate the trend in the mean number of categories per tool over time.

**Supplementary Figure 3:**
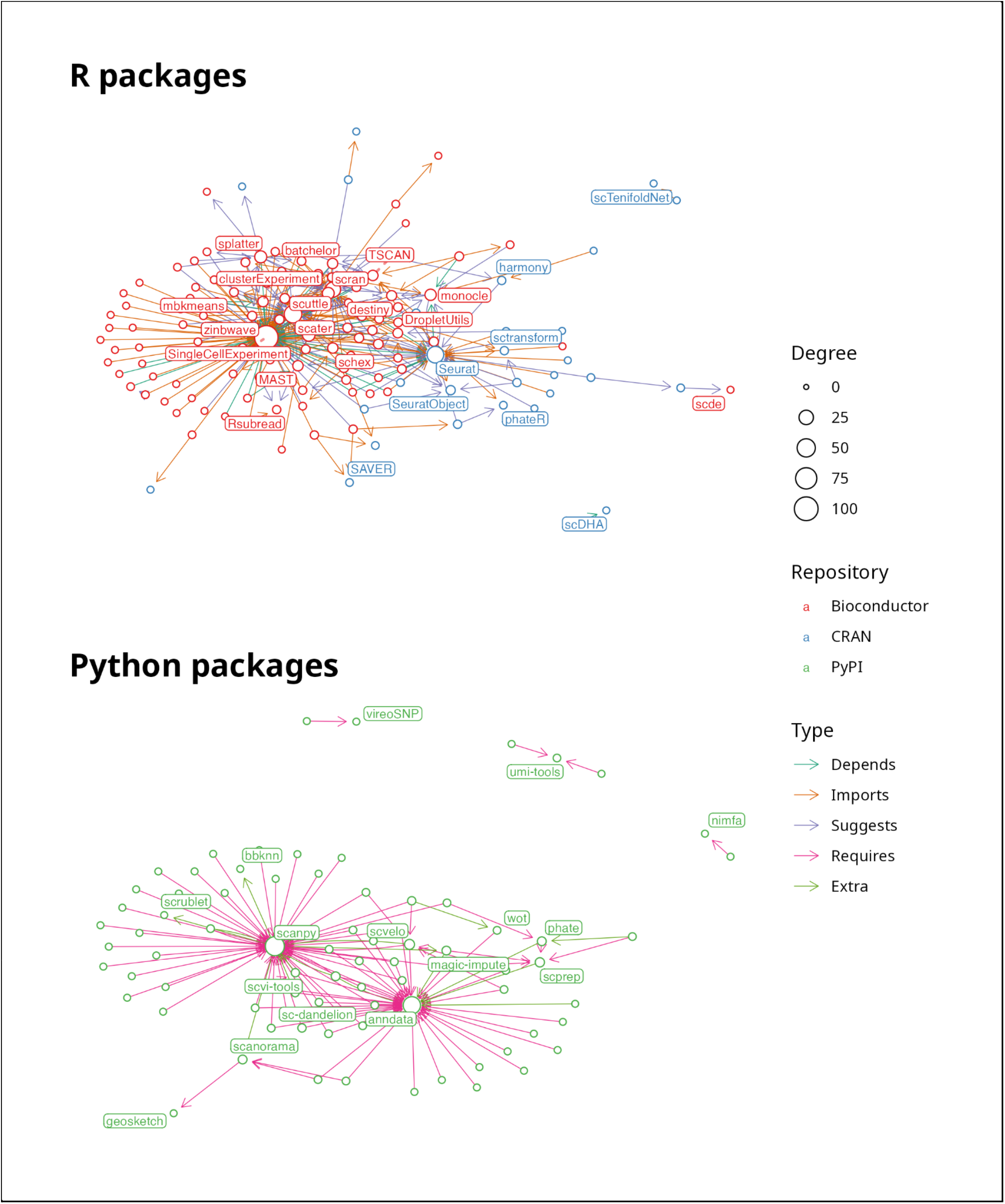
Dependency graphs for R and Python packages available from public software repositories. Nodes in the graph represent packages and edges represent dependencies. Node size shows degree and node colour indicates software repository. Edge colour indicates dependency type. Nodes with high PageRank centrality scores are labelled.

**Supplementary Table 1:** Coefficients and 95 percent confidence for publication models

| Term | Citations | Altmetric |
| --- | --- | --- |
| (Intercept) | -3.57 (-4.24, -2.90) | -0.25 (-0.98, 0.49) |
| log2(Num references + 1) | 0.34 (0.24, 0.45) | 0.37 (0.25, 0.49) |
| log2(Num authors) | 0.49 (0.36, 0.62) | 0.45 (0.32, 0.59) |
| Has preprint | 0.85 (0.60, 1.10) | 0.64 (0.39, 0.90) |
| Years (1st degree) | 6.27 (5.57, 6.98) | 0.66 (-0.07, 1.39) |
| Years (2nd degree) | 12.27 (11.11, 13.43) | 3.87 (2.68, 5.05) |
| Years (3rd degree) | 9.55 (7.79, 11.31) | 4.05 (2.26, 5.84) |

**Supplementary Table 2:**
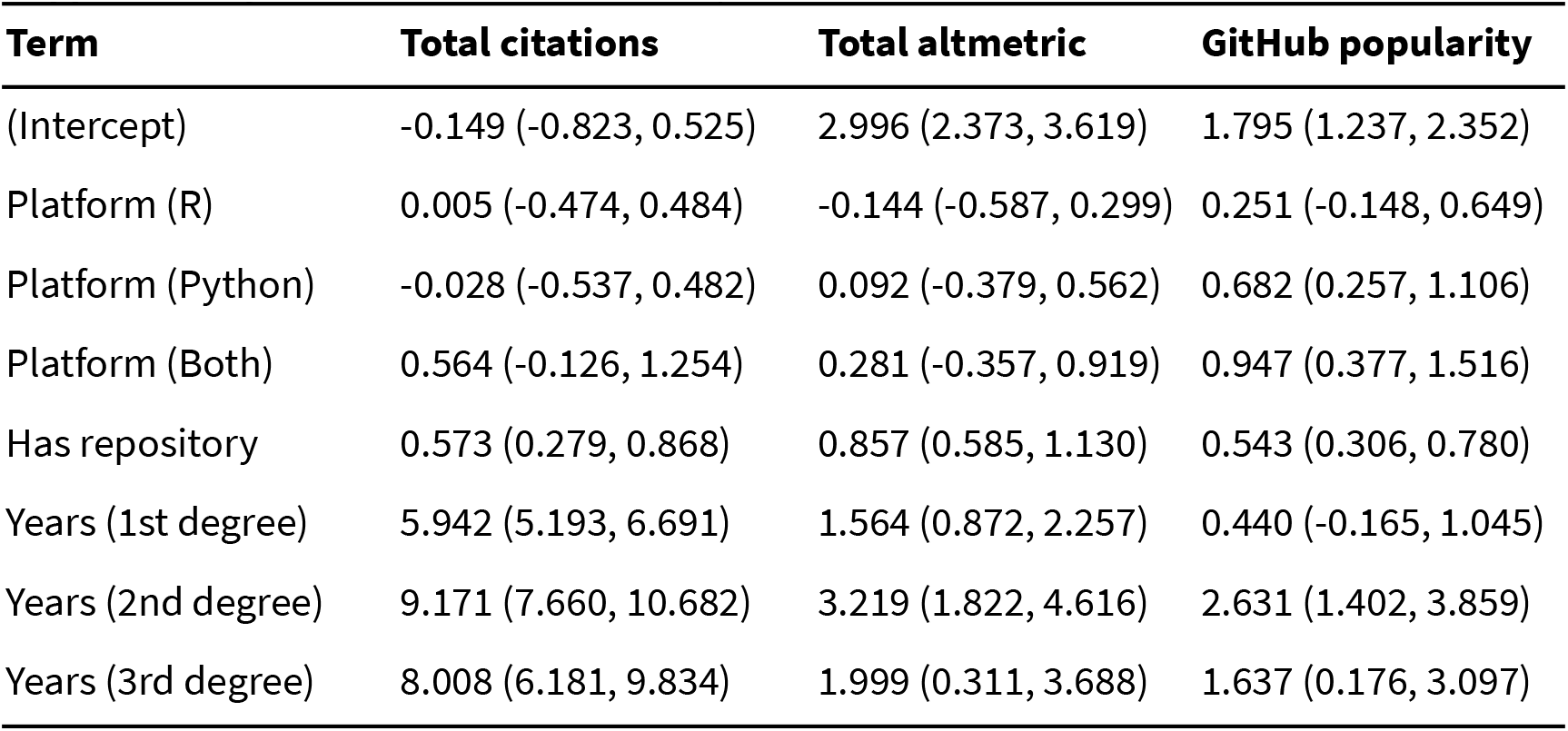
Coefficients and 95 percent confidence for tools models

| Term | Total citations | Total altmetric | GitHub popularity |
| --- | --- | --- | --- |
| (Intercept) | -0.149 (-0.823, 0.525) | 2.996 (2.373, 3.619) | 1.795 (1.237, 2.352) |
| Platform (R) | 0.005 (-0.474, 0.484) | -0.144 (-0.587, 0.299) | 0.251 (-0.148, 0.649) |
| Platform (Python) | -0.028 (-0.537, 0.482) | 0.092 (-0.379, 0.562) | 0.682 (0.257, 1.106) |
| Platform (Both) | 0.564 (-0.126, 1.254) | 0.281 (-0.357, 0.919) | 0.947 (0.377, 1.516) |
| Has repository | 0.573 (0.279, 0.868) | 0.857 (0.585, 1.130) | 0.543 (0.306, 0.780) |
| Years (1st degree) | 5.942 (5.193, 6.691) | 1.564 (0.872, 2.257) | 0.440 (-0.165, 1.045) |
| Years (2nd degree) | 9.171 (7.660, 10.682) | 3.219 (1.822, 4.616) | 2.631 (1.402, 3.859) |
| Years (3rd degree) | 8.008 (6.181, 9.834) | 1.999 (0.311, 3.688) | 1.637 (0.176, 3.097) |

## Notes

https://github.com/scRNA-tools/1000-tools-paper

### Summary of Updates

Fixed small typo in scTenifoldKnK tool name

